# Parallel high-performance computing algorithm to generate FEM-compliant volumetric mesh representations of biomolecules at atomic scale

**DOI:** 10.1101/578658

**Authors:** Jorge López, Salvador Botello, Rafael Herrera, Mauricio Carrillo-Tripp

## Abstract

The computational study of biomolecules has been undermined by the lack of models that accurately represent the structure of big complexes at the atomic level. In this work, we report the development of an algorithm to generate a volumetric mesh of a biomolecule, of any size and shape, based on its atomic structure. Our mesh generation tool leverages the octree algorithm properties with parallel high-performance computing techniques to produce a discretized hexahedral model faster than previous methods. The reported algorithm is memory efficient and generates volumetric meshes suitable to be used directly in Finite Element Analysis. We tested the algorithm by producing mesh models of different biomolecule types and complex size, and also performed numerical simulations for the largest case. The Finite Element results show that our mesh models reproduce experimental data.

## 1 Introduction

A derivation of the central tenet of Molecular Biology states that the structure of a biomolecule determines its function. Many syndromes and diseases are caused by the incorrect structuring of a certain biomolecule. Hence, efforts have been made to study the structure of biomolecules, their physicochemical properties, and the effect of changes in their structure on their function. In particular, there are many studies based on computational methods that try to explain the molecular mechanisms governing cellular processes. Even though there has been progress in our understanding of how these biological systems work, current molecular *all-atom* models have come short when the studied system is of considerable size (> 10^5^ atoms).

Molecular *coarse-grain* models were introduced to allow the study of larger complexes (a group of two or more associated biomolecules). Nevertheless, these models also have upper limits given by the current computational technologies. It is desirable to come up with a computational model that, keeping an atomic representation, can describe very large biological systems, e.g., a virus, a cellular organelle (a specialized subunit within a cell that has a specific function), or even the whole cell.

Volumetric meshes are a potential candidate to overcome such a problem. Not only do they give an atomic-level description of a large system, but they are also compatible with numerical methods (e.g. the Finite Element Method or FEM) in order to perform simulations of a certain biophysical process. However, the generation of a volumetric mesh of *good quality* is not an easy task. Several factors have to be taken into account, namely, shape of the mesh elements, their size distribution, regularity, etc.

In this work, we describe the development of an algorithm designed to discretize the volume occupied by all the atoms of a given biomolecule or biocomplex. The discretization process generates an approximation to the original biosystem through space decomposition combined with parallel and high-performance computing techniques. This approach makes efficient use of memory, and the multi-platform multiprocessing implementation reduces the execution time by several orders of magnitude in comparison to previous developments. The output is a volumetric mesh suitable for FEM analysis.

We applied the algorithm to generate a volumetric mesh of representative examples of all major categories of molecules relevant to biological systems. These categories are *carbohydrates*, *lipids*, *nucleic acids*, and *amino acids*. The specific systems chosen for our study are described in Table 1 and illustrated in Figure 1. Given its importance in life as an organic solvent, bulk water in liquid state was also included. In this study, we have been careful to consider a large diversity of biomolecules regarding they size, shape, and function.

**Table 1.**
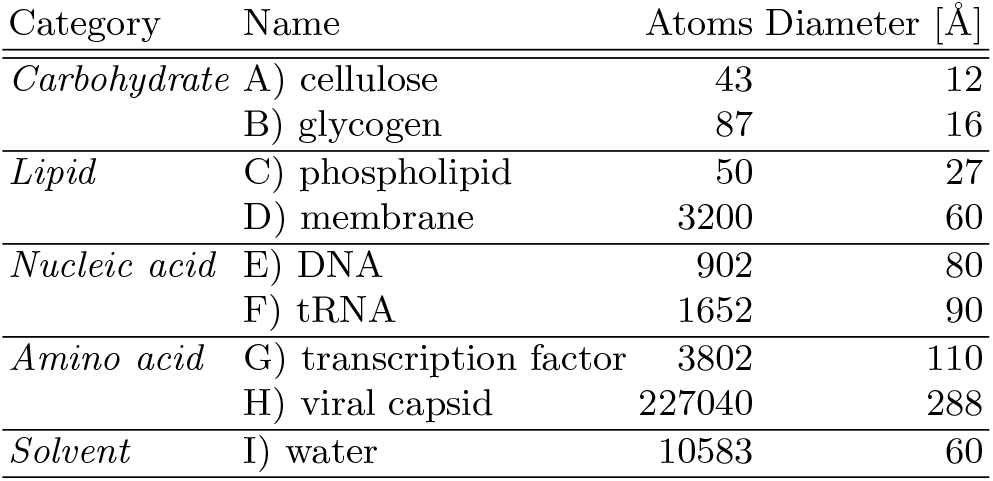
Major categories of biomolecules considered in this study. Two representative systems were chosen in each case, either a single molecule or a complex, plus water. Number of atoms and characteristic size are shown.

| Category | Name | Atoms | Diameter [$\text{\AA}$] |
| --- | --- | --- | --- |
| *Carbohydrate* | A) cellulose | 43 | 12 |
|  | B) glycogen | 87 | 16 |
| *Lipid* | C) phospholipid | 50 | 27 |
|  | D) membrane | 3200 | 60 |
| *Nucleic acid* | E) DNA | 902 | 80 |
|  | F) tRNA | 1652 | 90 |
| *Amino acid* | G) transcription factor | 3802 | 110 |
|  | H) viral capsid | 227040 | 288 |
| *Solvent* | I) water | 10583 | 60 |

**Fig. 1.**
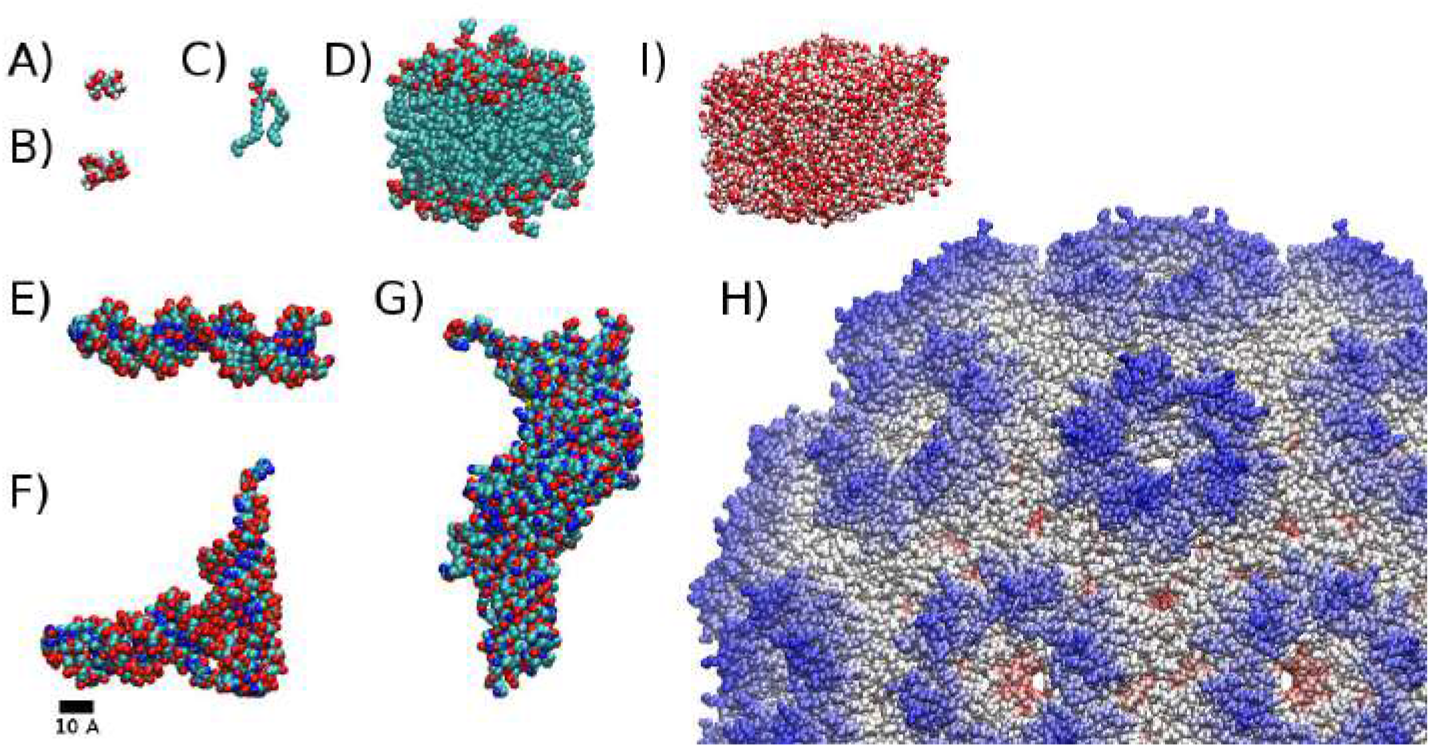
Biomolecules considered in this study: A) cellulose, B) glycogen, C) phospholipid, D) small section of a biomembrane (64 phospholipids), E) DNA (22 nt long), F) tRNA, G) transcription factor, H) icosahedral viral capsid, I) 1×10^−19^ ml of bulk liquid water. In all the cases, except for the viral capsid, colors correspond to the element type: Carbon in cyan, Hydrogen in white, Oxygen in red, and Nitrogen in blue.

Furthermore, in order to test the compatibility of the mesh generator with FEM, we carried out numerical simulations of nanoindentation on the viral capsid. There are experimental results of Atomic Force Microscopy (AFM) where the authors *squeeze* the capsid while measuring the applied force and the produced deformation on the complex structure [Michel2006,Arkhipov2009,Roos2010]. We estimated the Young’s modulus for the capsid by using the spring constant derived from the experimental force-indentation profile in the linear regime. We show that our model is able to reproduce experimental results and is in agreement with previous models.

## 2 Meshing

The state of the art in meshing techniques-methods can be summarized in three categories: Advancing Front (ADF), Delaunay Triangulation (DT), and Space Decomposition (SD). Each meshing technique presents advantages and drawbacks, so it is important to choose the correct technique in order to produce a mesh compliant for an accurate and fast FEM analysis. This work is focused on a particular SD method given by an octree algorithm due to its advantages in managing computational and memory complexity.

### 2.1 Delaunay Triangulation

This technique tackles the mesh generation problem by performing efficient geometric operations and has been widely studied [Paul1990], [Pascal2008], [Siu2013], [Paul1998], which has improved its computation time. DT methods are highly regarded by researchers and engineers since numerical methods for solving PDE models of real life problems require high quality triangulations.

DT generates good 2D meshes. However, it presents a significant drawback in 3D: it may generate elements with null volume which comply with the Delaunay properties. Some implementations tackle this problem by relaxing the Delaunay condition [Paul1990].

DT takes a set of points ℙ and creates a triangulated mesh called the convex hull [Pascal2008], [Siu2013]. This mesh can be used to perform numerical simulations, but if some elements have volume nearly zero, the stability and accuracy of the numerical solution may be affected. Thus, some modifications may be needed, such as removing collapsed elements and refining the mesh to improve the mesh quality.

The method’s main advantages are its high speed and robustness. On the other hand, the main drawback is that it does not preserve the meshing domain. Other algorithms [Weatherill1992] can be applied to circumvent this problem but with the cost of losing efficiency and increasing computation time. Finally, this technique is difficult to be parallelized since the creation of elements requires the information of all the triangles contained in its circumscribing circle, causing concurrency problems. Nevertheless, some parallel implementations can be found in [Hardwick1997] and [Cigoni1993].

### 2.2 Advancing Front

This approach starts on the boundary and inserts new points inside the domain. These points are used to create triangles (2D) or tetrahedrons (3D), but they can be also used to create quadrilateral or hexahedral elements by applying slight modifications.

A well-known mesher based on ADF is NETGEN [Schöber1997]. It is customizable in the sense that the user sets parameters to determine the mesh size and the optimization steps to apply on the generated mesh.

The ADF technique requires more time than the comparing techniques, due to its complexity. It could be said that this mesh generation technique generates triangles nearest to equilateral due to the tests performed before it creates a new element. This technique is based on local operations and, as a consequence, it is highly parallelizable.

### 2.3 Space Decomposition

SD methods are equipped with features for creating meshes with acceptable quality in a short time, using highly parallelizable methods. Thus, this methods are, possibly, the most widely used by FEM researchers.

Broadly speaking, there exist two types of SD methods to create FEM-suitable meshes. The first one is *the bin* method, and the second one is the *octree* technique. They are based on performing bisections over each dimension depending on a maximum level of refinement determined by the user. Although both algorithms look similar, the octree technique performs bisections based on the domain to be meshed, while the bin technique divides the bounding box on cells of the same size over each dimension, which causes excessive memory requirements. In octree, apart from the low usage of memory, cells can be represented as binary numbers, which allows us to move through the octree by using bitwise operations, thus accelerating search operations in contrast with other techniques.

The octree algorithm can be used to easily generate tetrahedral or hexahedral meshes, unlike other techniques that need to be modified in order to create hexahedral meshes.

One of the main drawbacks of SD techniques is their inability to preserve corners, usually known as sharp features, since the octree is based on quadrilateral elements complex domains cannot be well-fitted with these geometric forms. The problem can be tackled by performing some modifications on boundary elements in order to achieve the best possible approximation to the original domain.

The main advantage of this method is that meshes can be created for most domains due to the spacial adaptability of the method. Another advantage is the possibility of having efficient parallel implementations which lead to fast generation of high quality meshes. In the implementation, all the elements are equal, which will lead to improvements in the numerical solutions.

#### Remark

The three techniques share some features in the sense that they perform similar operations. For instance, DT and SD require a ray casting technique that can be carried out using computer graphic strategies and can lead to create an inefficient algorithm. However, since the octree nodes are aligned, the ray casting technique can be implemented taking advantage of its structure. In summary, the three meshing techniques need to perform geometric tests in order to create the best possible triangulation. While SD methods have control over all the objects contained in the working space so that the number of geometric tests is reduced due its local operation, the DT and ADF methods have difficulties controlling objects inside the domain.

## 3 Octree mesher

The octree is an SD method and it must be used in a delimited space. The overall efficiency of the algorithm can be increased considerably when the space is defined correctly. The best option is to work on a normalized domain such as the square [0, 1] × [0, 1] in 2D, or the cube [0, 1] × [0, 1] × [0, 1] in 3D.

The first step is to scale the object *Ω* we wish to mesh to fit into the square or the cube, whose boundary is the bounding box (see Figure 2). The next step is to subdivide the box progressively in order to choose only the smaller boxes (octree cells) that intersect *Ω*, and thus obtain a volume made up of small octree cells that closely approximates *Ω*.

**Fig. 2.**
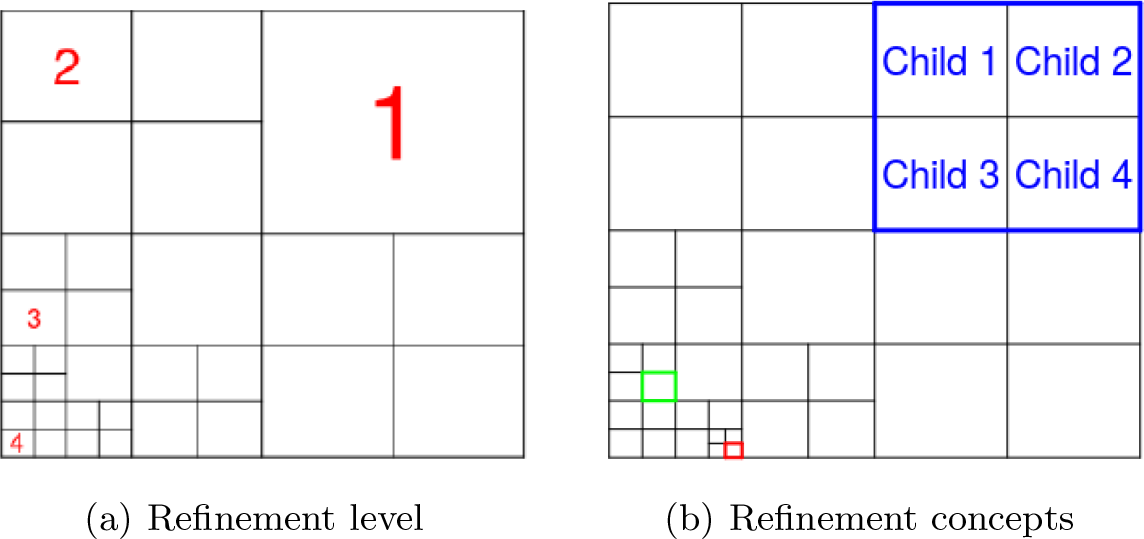
Refinements generalities

The subdivision generates children from a given parent cell (see Figure 2(b)). Cells to be bisectioned are selected based on intersection tests. The traditional octree method performs intersection tests between cells and triangles, and the way to perform it efficiently is by using computer graphics strategies, which can help to reduce considerably the testing time.

The octree subdivision generates octree cells with different sizes that are associated to the concept of *refinement level*. In Figure 2(a) there are red numbers indicating the refinement level of each cell. The bounding box is considered as a zero refinement cell.

The octree can be used either to generate hexahedral or tetrahedral meshes. In order to create the tetrahedral meshes, it is required to accomplish the constrain 2:1, i.e. for each cell, all its neighbors in all directions must have a difference of at most one level of refinement. For example, the green square in Figure 2(b) shows a cell which fulfills the constraint. On the other hand, the red cell does not fulfills the constraint because the right neighbor has two refinements levels below the tested cell. However, in our implementation, the constrain is not required because the hexahedral meshes generated have all its elements of the same size.

The main purpose of scaling *Ω* to the normalized square or box to encode the order of the octree cells in terms of binary numbers and to be able to move through the complete octree using bitwise operations that are faster than traditional arithmetic operations. This is, in fact, the main HPC feature of octree that reduces computing time, although its implementation may be challenging. The binary implementation to manage octree cells is based on [Frisken2002].

Figure 3 shows the way to work with the bisections and binary coding using a simplification to 1D. First image 3(a) is the root or initial segment of line. This line lies in the domain [0, 1] and has the binary coding shown on top. The first bi-section generates Figure 3(b) where two segments have been created and two binary numbers represent them. Figure 3(c) shows one more partition, where we see four segments and their binary coding. Finally, adding two more bisections as in Figure 3(d), there are six line segments, each one with a binary number associated to it (blue numbers). It is easy to convert from binary to decimal by using the line segment refinement level and binary codding.

**Fig. 3.**
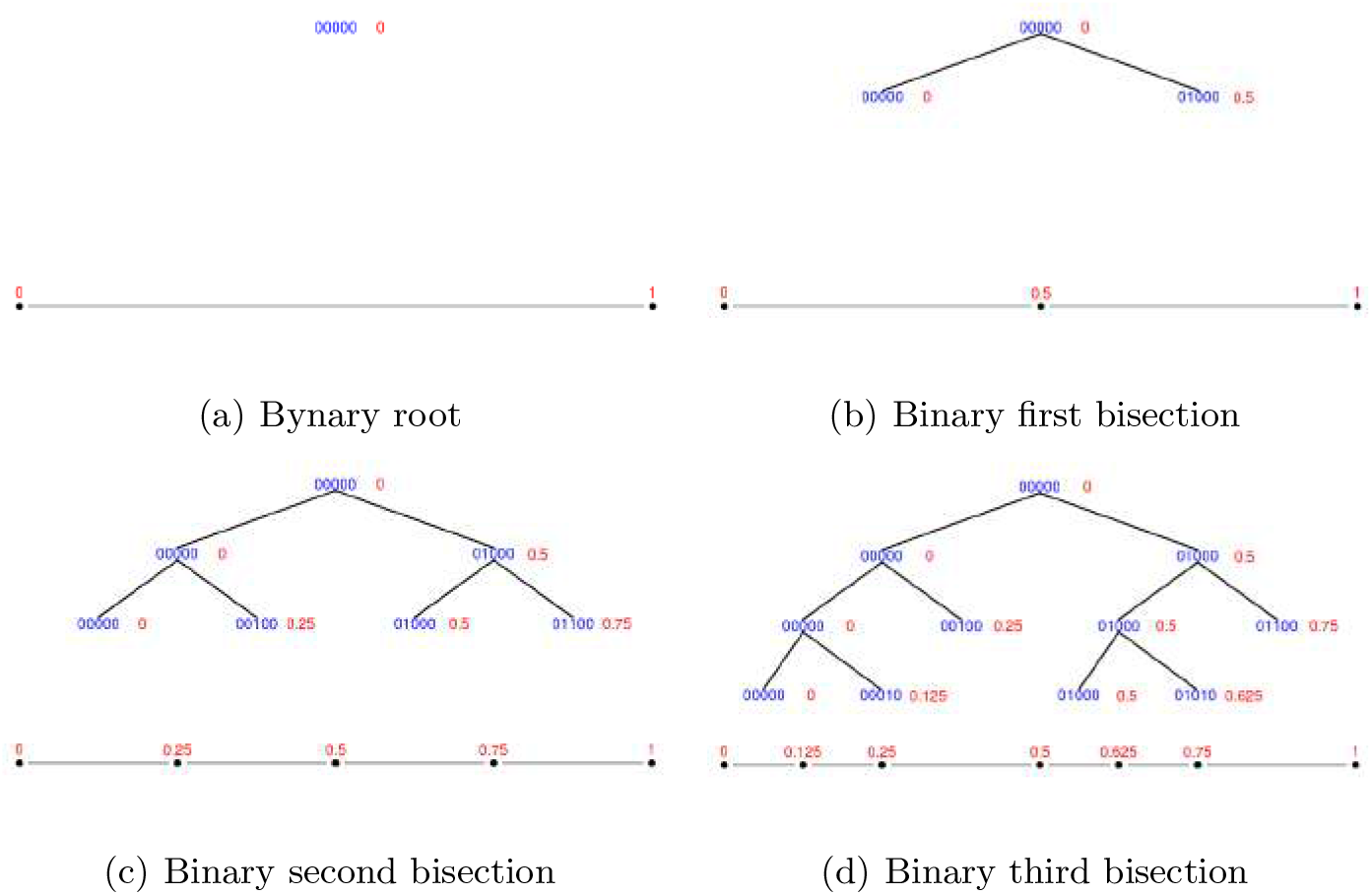
Binary codification on octree

The selection of cells belonging to the final mesh is naturally achieved during subdivision of the octree, keeping the elements that intersect *Ω*. The octree cells that become part of the final mesh must fulfill the following features:

– Two distinct elements must not overlap.
– In 2D, an edge is only shared by two adjacent elements; in 3D, a face is only shared by two adjacent elements.
– The elements must to be positively oriented (correct node labeling in FEM).

The final step is to return *Ω* to its original scale together with the selected octree cells.

The time reduction when using the octree mesher is achieved by using parallel computing in some of the operations, as well as the use of some HPC techniques (binary codification). Using this parallel scheme, meshes are generated in one computer by using, concurrently, several cores. However, if the computer has a processor with more than one core and all the cores share the same RAM, all of them can modify the same variables at the same time, which is to be avoided since it could lead to erroneous results or low parallel performance. Nevertheless, OpenMP is suited for high-level parallelization in a simple manner with a low number of prepocessor instructions (shared memory parallel scheme [OpenMP2018]). Our main objective is not only to parallelize the implementation but to do so in order to achieve a highly efficient parallelization. Hence, we must take care of concurrent operations and data synchronizations. Some of the parallel operations in the proposal are:

– **Octree refinement:** Since each and every cell is independent, various cells can be refined at the same time by using as many cores as possible.
– **Cells nodes setting:** This operation adds all the coordinates to the octree cells. Similarly, this operation is parallelized due to the independence of the information added to a cell.
– **Intersection tests:** The independence of the cells means that various cells can be tested for intersection with spheres at the same time.

### 3.1 Intersection test for meshing

In general, biomolecules are represented by clustered sets of spheres (one for each atom present). Thus, the intersection test of the octree method must be modified slightly. Because we only process the spheres contained in the cell that is being subdivided, the total number of intersection tests is reduced.

One more advantage of our proposal for meshing biomolecules is that all the elements in the resulting mesh are equal. This implies that the solution of any structural analysis through numerical methods will be carried out in an efficient way, saving memory and computing time.

### 3.2 Biomolecules

Our meshing strategy can process biomolecules of any size and shape, unlike previous methodologies. Hence, it can generate a mesh for non-symmetrical structures. An example of the progress in atomic detail achieved by increasing the resolution value is shown in Figure 4 for the cellulose molecule. Lower values give higher resolution. We generated a mesh for all the data set described earlier. Results are shown in Figure 5 and their characteristics are given in Table 2. In this case, the mesh resolution is different for each biomolecule due to their different sizes.

**Fig. 4.**
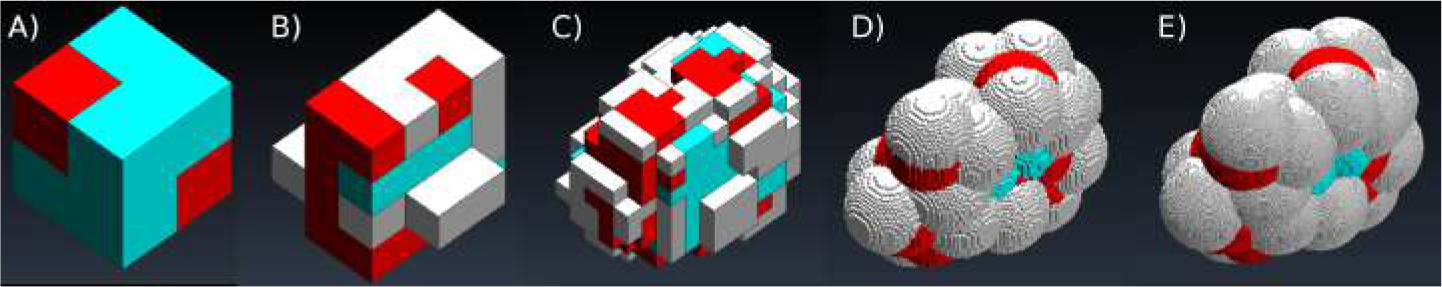
Cellulose octree mesh generated using different resolutions: A) 6.0, B) 3.0, C) 1.0, D) 0.1, E) 0.03 Å. The colors correspond to the element type: Carbon in cyan, Hydrogen in white, and Oxygen in red.

**Fig. 5.**
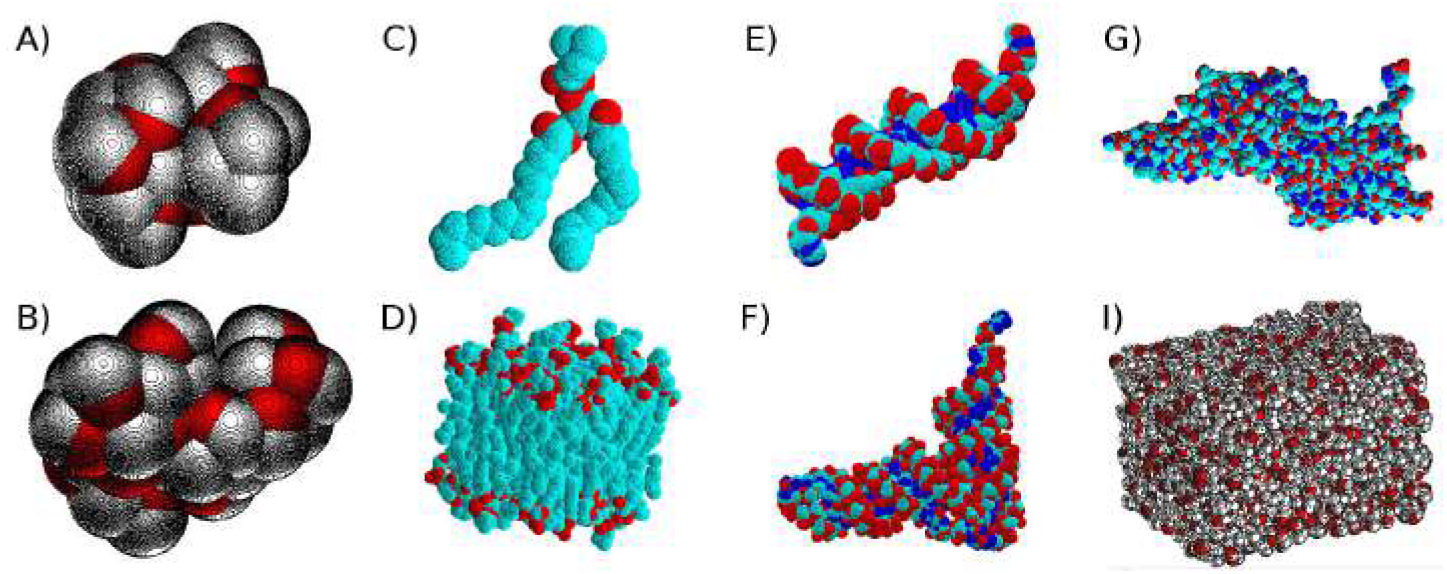
OctreeMesh representation of biomolecules considered in this study. A) Cellulose, B) glycogen, C) phospholipid, D) small section of a biomembrane (64 phospholipids), E) DNA (22 nt long), F) tRNA, G) transcription factor, I) 1×10^−19^ ml of bulk liquid water. The colors correspond to the element type: Carbon in cyan, Hydrogen in white, Oxygen in red, and Nitrogen in blue.

**Table 2.** OctreeMesh features for the biomolecules considered in this study. Mesh resolution given in Å.

| Category | Name | Mesh | Mesh | Time [s] |
| --- | --- | --- | --- | --- |
|  |  | Resolution | Elements |  |
| *Carbohydrate* | A) cellulose | 0.0375 | 8,862,661 | 200.92 |
|  | B) glycogen | 0.0450 | 9,069,479 | 220.12 |
| *Lipid* | C) phospholipid | 0.0460 | 4,672,597 | 106.06 |
|  | D) membrane | 0.2250 | 3,934,749 | 95.74 |
| *Nucleic acid* | E) DNA | 0.1300 | 8,790,359 | 246.40 |
|  | F) tRNA | 0.1530 | 4,973,299 | 122.32 |
| *Amino acid* | G) transcription factor | 0.2100 | 5,272,603 | 133.26 |
|  | H) viral capsid | 5.0000 | 4,709,364 | 114.83 |
| *Solvent* | I) water | 0.2400 | 6,405,468 | 160.16 |

## 4 Viral capsids

The protein shell encapsulating the genome material of a virus is known as the capsid. Its fundamental functions are to protect and transport the viral genome, as well as helping recognize the host cell. Detailed knowledge about the physical properties that characterize the viral capsid have proved to be very important in structural biology, biotechnology, and medicine. On the one hand, the analysis of virus nanoindentation by AFM has been used as the standard way to study the mechanical response of capsids as well as understand their molecular structure. On the other hand, numerical simulations of the nanoindentation process have allowed the estimation of physical parameters such as the Young’s modulus. Various descriptions of this type of macromolecular complex have been proposed at different scales for their study.

### 4.1 Capsid meshers

Only a few works address the problem of meshing a capsid to be used in numerical simulations, mainly due to their large size and geometrical complexity. In [Sanner1996], the authors present a surface meshing algorithm for the molecular complex. This algorithm was one of the first to avoid the self intersecting problem when the surface mesh was created. In [Yu2008], the authors introduced a new approach that could generate the surface and the volumentric mesh of a capsid, tackling for the first time “feature preservation”. This algorithm generated quality meshes in a considerable amount of time. In [Cheng2009], the DT strategy was used to create surface meshes with thousands of elements in minutes. A recent proposal was focused on volumetric mesh generation, as opposed to surface mesh generation [Alonzo2018]. A volumetric mesh of the capsid was created using symmetry features, meshing a geometric unit and rotating it to generate the full capsid. The main drawback of this method is the need to delete repeated nodes, which implies a computational cost increase for the creation of high resolution meshes. In that sense, the best feature of our octree proposal is the comparatively very short computing time needed to generate the capsid mesh.

### 4.2 Nanoindentation

In an AFM experiment, the relation between the magnitude of an applied force and the capsid’s indentation (biocomplex deformation) can be measured. As a result, a characteristic force-indentation graph is recorded. The slope of the curve in the linear regime is related to the capsid’s spring constant.

The numerical simulation of an AFM experiment (Fig. 6) consists in aligning a particular capsid’s symmetry axis with an applied force load. Such force is directed over the top of the mesh, while the bottom is fixed on a base. We used the FEMT open source code [Vargas2012] to carry out the numerical simulation. In this work, we carried out numerical simulations of the nanoindentation of a whole viral capsid using volumetric meshes generated by the octree space decomposition method. The quality of the meshes produced by the algorithm is validated by reproducing the nanoindentation simulations results presented in [Alonzo2018] for the CCMV.

**Fig. 6.**
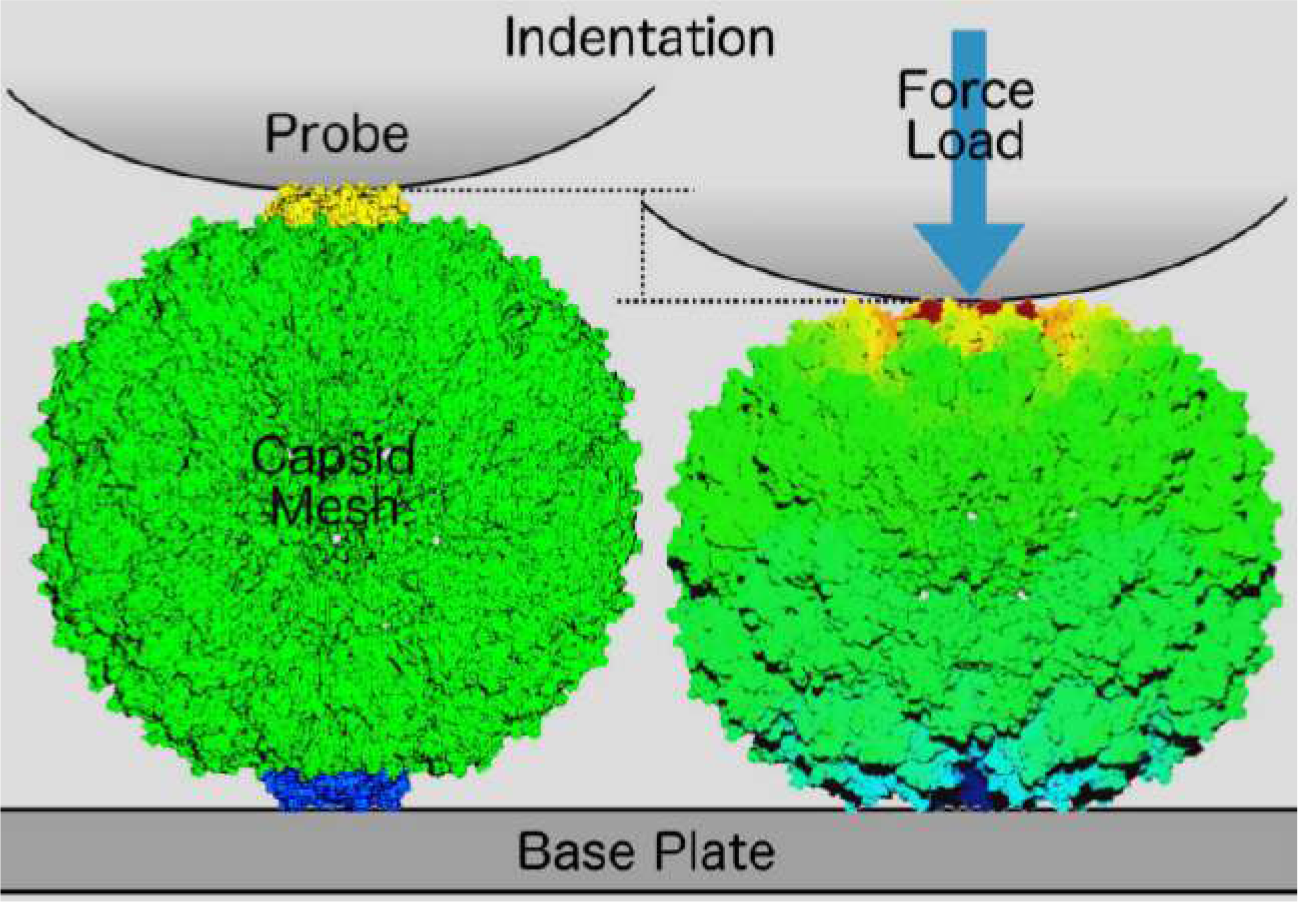
Capsid nanoindentation simulation. Left: Initial conditions, where the mesh elements in blue are in direct contact to a base, and mesh elements in yellow are in direct contact with a probe. Right: A force load is applied on a given capsid’s symmetry axis. The mesh deformation is quantified by the indentation produced (displacement measured in each mesh element). The color scale used represents minimum displacement in blue and maximum displacement in red.

A summary of the results are shown in Table 3. The first column indicates the virus that the capsid belongs to, *T* indicates the geometrical protein arrangement, *d* is the biocomplex diameter (2 times the outer radius), *w* is the capsid width (outer radius minus inner radius), nanoindentation *I* (16% of *d*), force needed to produce *I* on the 5-fold axis, Young’s modulus *E* prediction, and experimental spring constant *k*_*exp*_ [Michel2006].

**Table 3.** Nanoindentation of a wild type empty capsid and mechanical properties of Cowpea chlorotic mottle virus (CCMV), using a mesh resolution of 1 Å. Shown: geometrical protein arrangement *T*, diameter *d*, capsid width *w*, nanoindentation *I*, force needed to produce *I* on the 5-fold axis, Young’s modulus *E* prediction, and experimental spring constant *k*_*exp*_ [Michel2006].

| Capsid | T | d | w | I | F | E | $k_{exp}$ |
| --- | --- | --- | --- | --- | --- | --- | --- |
| Name | Number | Å | Å | Å | nN | MPa | N/m |
| CCMV | 3 | 288 | 50 | 46 | 2.24 | 193 | 0.15 |

A comparison of the structural analysis results between a previous algorithm, CapsidMesh [Alonzo2018], and the octree algorithm are shown in Figure 7. The maximum displacement shows that both algorithms produce indentations almost identical. The stress analysis shows some differences in intermediate resolution values (2-6 Å), probably due to the shape of the mesh elements and the difference in the cavity shapes that are naturally present with these topologically faithful meshing methods. More importantly, a significant difference is observed between the two algorithms in terms of mesh generation times. The CapsidMesh processing time tends to increase exponentially when the mesh resolution is increased (lower values). Although the octree computing time also increases, the efficient and parallel implementation plays an important role in saving time. For instance, it takes 261 seconds to generate a mesh with resolution 1Å, while CapsidMesh needed 85,598 seconds. The structural results were mapped into the capsid’s mesh representation to illustrate the location of interesting features, shown in Figure 8.

**Fig. 7.**
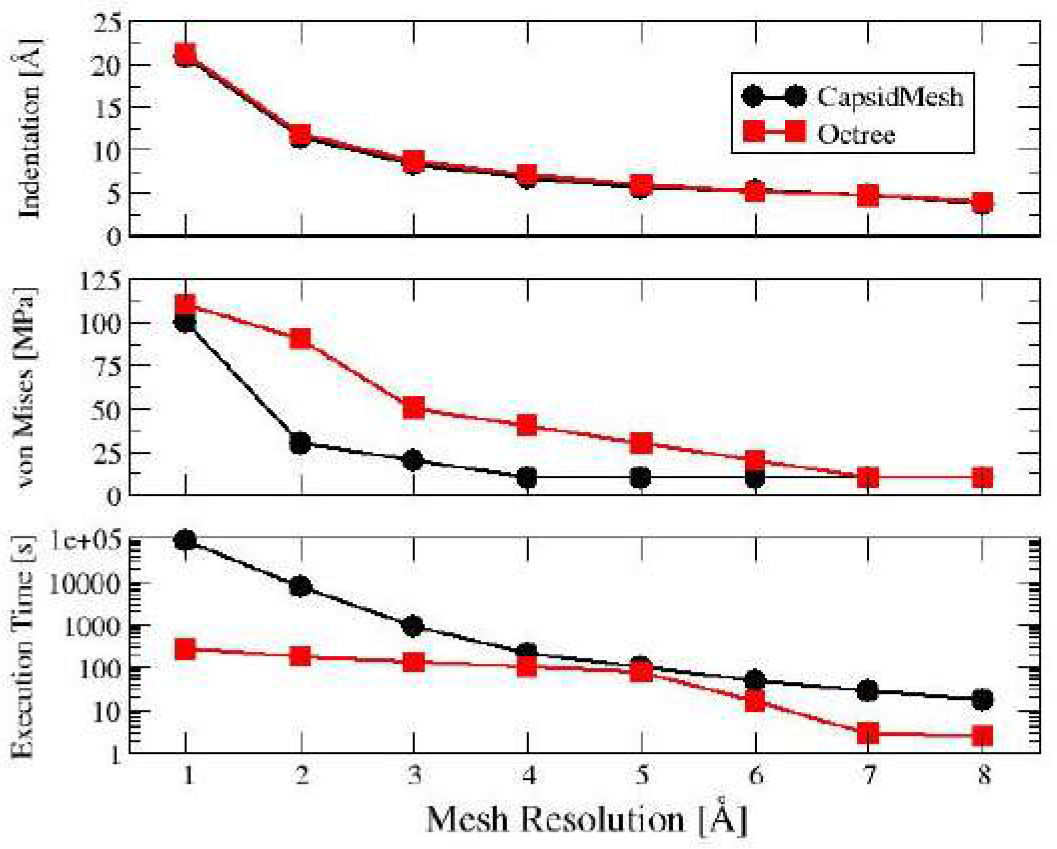
Capsid meshing and nanoindentation results, comparison to previous method. From top to bottom: Indentation produced, structural stress analysis, and time spent to produce the mesh, as a function of mesh resolution.

**Fig. 8.**
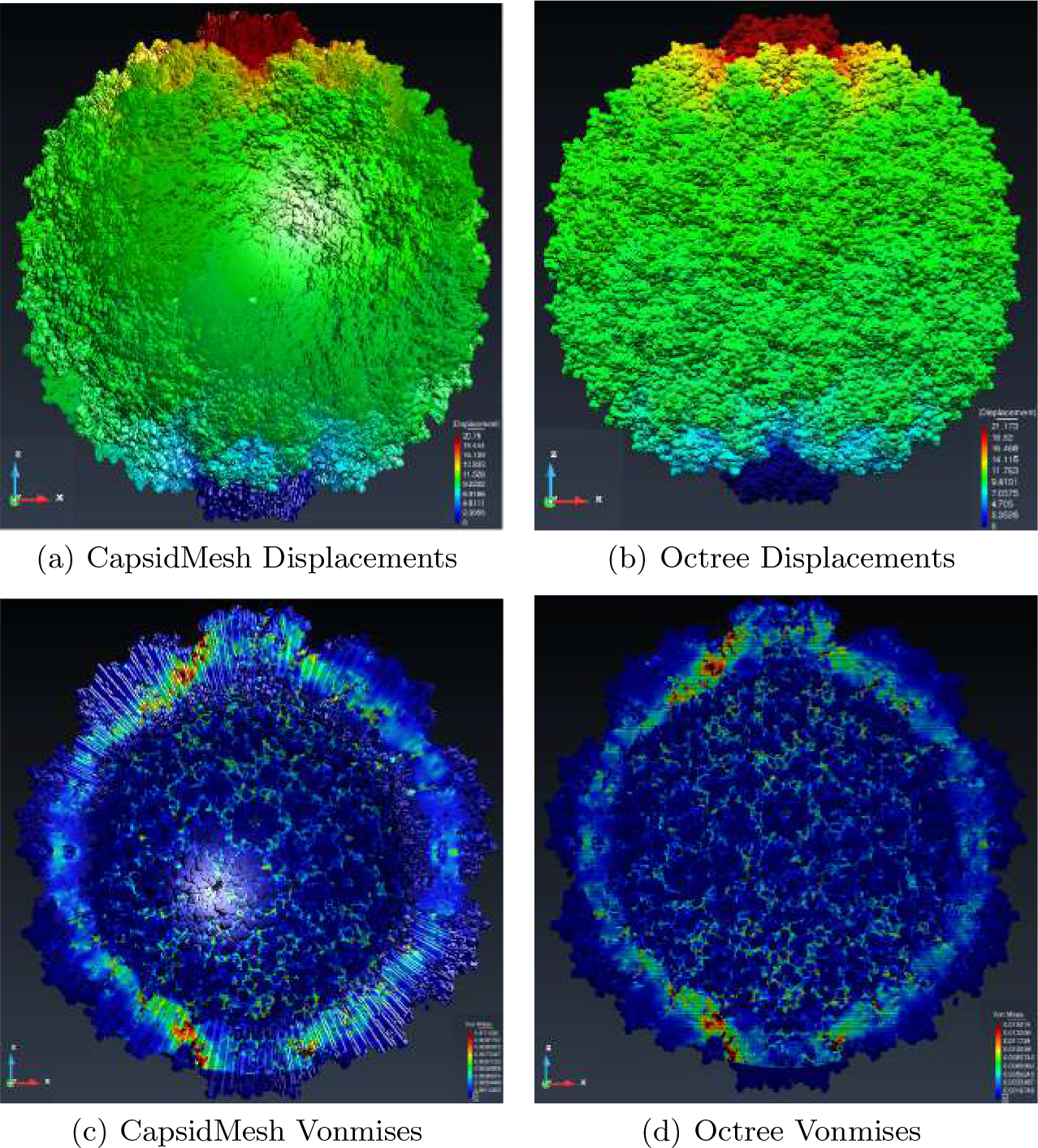
Simulation results

## 5 Conclusion

In this work, we report the development of a computational algorithm to produce volumetric meshes of biomolecules based on the volume and position of their constituent atoms. Discretization of space is accomplished by the octree method. Parallelization techniques allow for an efficient use of memory and fast running times. Since the mesh elements have an hexahedral geometry, our algorithm can generate a volumetric mesh of any biomolecule or biocomplex, independent of its size or shape.

We built mesh models of representative examples of the four major categories of biomolecules, spanning a large diversity of biological structures and functions. An important feature of these meshes is that they are FEM-compliant, i.e., they can be directly used to perform numerical simulations of a given biophysical process. To test this, we carried out the simulation of the nanoindentation of a full viral capsid represented by our mesh. This is the largest biocomplex in our data set.

Results show that our algorithm can be used to model very large biocomplexes, even with more than 200,000 atoms, and still keep atomic scale. When executed in a single-core computer, the octree mesher can generate volumetric meshes of up to 1×10^7^ elements. This number increases to 1×10^8^ when the algorithm is executed in a multi-core computer or HPC cluster. Also, the mesh resolution can be higher than 0.5 Å. Compared to previous methods, the one reported here is orders of magnitude faster, faithfully reproduces the shape of the biological system, and has a stable behavior during numerical simulations.

